# A structural mechano-chemical model for dynamic instability of microtubule

**DOI:** 10.1101/291682

**Authors:** Shannon F. Stewman, Ao Ma

## Abstract

Microtubules are a major component of the cytoskeleton and vital to numerous cellular processes. The central dogma of microtubules is that all their functions are driven by dynamic instability; understanding its key phenomena (i.e. catastrophe, rescue, pause, differential behaviors at the plus and minus ends) distilled from a myriad of experiments under a consistent and unified scheme, however, has been unattainable. Here, we present a novel statistical-physics-based model uniquely constructed from conformational states deduced from existing tubulin structures, with transitions between them controlled by steric constraints and mechanical energy of the microtubule lattice. This mechano-chemical model allows, for the first time, all the key phenomena of dynamic instability to be coherently reproduced by the corresponding kinetic simulations. Long-puzzling phenomena, such as aging, small GTP-cap size, fast catastrophe upon dilution and temperature-induced ribbon-to-tube transition of GMPCPP-tubulins, robustly emerge and thus can be understood with confidence.

## I. Introduction

Microtubules (MT) are crucial for numerous essential cellular processes, including mitosis, morphogenesis, and neurogenesis [1–5]. All functionality of MTs is driven by dynamic instability, which has long been characterized and quantified. But despite intensive experimental and modeling efforts of three decades, understanding all the key phenomena of dynamic instability within a unified and consistent scheme has been unattainable [1, 6]. This has been the bottleneck to further understanding the cellular MT network because reductionist attempts in this regard had to rely on empirical notions of how regulatory proteins affect the phenomena rather than the mechanism of dynamic instability, leading to puzzling and inconsistent results [1].

Microtubules are cylindrical polymers of α,β-tubulin heterodimers arranged head-to-tail in thirteen protofilaments (PFs), with the plus and minus ends marked by the β- and α-tubulin, respectively. Dynamic instability refers to the spontaneous stochastic switching of a MT end between steady-states of growth and shortening [2, 7], manifesting four key phenomena: 1) catastrophe (growth to shortening), 2) rescue (shortening to growth), 3) a meta-stable pause state (neither growth nor shortening) [2, 8], and 4) the plus and minus ends both show dynamic instability, but with very different behaviors [8]. Among these four phenomena, efforts since 1984 have focused on plus-end catastrophe alone, yet its mechanism remains unclear [1, 2, 6, 7, 9].

The prevalent idea on plus-end catastrophe has been that a structure at the growing end caps a MT from depolymerization, with two hypotheses: the GTP-cap and the structural cap [7, 9, 10). The GTP-cap hypothesis assumes that GTP-tubulins adopt a straight conformation characteristic of stable MTs, and GDP-tubulins prefer the curved form during rapid shortening. Thus MTs comprise mechanically-strained GDP-tubulins held straight by a cap of GTP-tubulins at the growing plus-end. The cap disappears when hydrolysis catches up with the growing tip, and depolymerization ensues [2, 7]. The structural cap hypothesis derives from the observation that a plus-end grows as either a tube or a curved sheet [11]. The curved sheet was postulated as a structural cap that stochastically closes into a tube, and catastrophe follows full tube closure.

Experiments to determine the size and nature of these caps have led to conflicting conclusions [12–17]. The finding that even long MTs have a small GTP-cap of one to three rows [12] was puzzling because the mechanical strain in the lattice that eventually causes catastrophe should increase with MT length [13, 14], although this finding has been challenged from a different perspective recently [15]. Moreover, Gardner et al showed that catastrophe is multi-step [16], contrary to the long-held belief that it is first-order but corroborating an earlier proposition by Odde et al [17]. Structural studies, on the other hand, suggested tight coupling between dynamic instability and the structures of tubulins [11, 18–21), although its mechanism remains unclear.

Computational models have also been built upon the cap concepts to clarify details of the mechanism of plus-end catastrophe [6, 9, 22-31]. They typically have spatial patterns that signal catastrophe through the loss of the GTP-cap, and ad hoc rules for hydrolysis [6, 23–31). These models suffered conceptual difficulties because they placed the critical step of catastrophe on hydrolysis, which is an intrinsically local event dictated by the inter-dimer longitudinal interface housing the GTP [18, 19], whereas catastrophe is a collective emergent phenomenon unique to MTs (*vide infra*), implied in its being a rare-event orders of magnitude slower than growth.

Consequently, catastrophe in these models often appears sensitive to the specific GTP-cap patterns that are difficult to interpret physically [6], and the ad hoc rules for hydrolysis, which effectively correlate hydrolysis at different sites to create many-body events, contradict our understanding of chemical reactions [6, 29, 30]. In addition, there were studies focusing on unifying the non-equilibrium assembly dynamics of microtubule and actin from a general view. Zong et al studied stationary solutions of the master equations of a one-dimensional model of microtubule/actin using a novel variational method and analyzed the effects of different rate parameters on the properties of the steady states [32].

To date, all four key phenomena of dynamic instability remain puzzling: catastrophe at the plus-end is poorly understood despite efforts of three decades; rescue has seen little systematic investigations [33]; pausing and dynamic instability at the minus-end have not yet been explored. This situation has obstructed further understanding of the cellular MT network and urges a new systems induction approach to modeling dynamic instability. The correct model should comprehensively and consistently explain and reproduce all the phenomena of dynamic instability, as they all reflect the same underlying physical machinery. Consistency is vital because it constrains the model by eliminating alternative mechanisms that are plausible for some phenomena but contradicted by others. Comprehensiveness is critical because, given the enormous complexity of the MT system, focusing on only a subset of phenomena places inadequate constraints on the model, so it will inevitably be incompatible with the omitted phenomena, thus breaking the consistency principle.

Here we present a structural mechano-chemical model that, contrary to previous efforts that focused on plus-end catastrophe, unifies all phenomena of dynamic instability coherently and consistently. To our knowledge, no such model currently exists. According to our model, dynamic instability is driven by transitions among conformational states, which are deduced from existing tubulin structures [18, 19, 21] but have eluded previous investigations. Steric constraints on tubulin structures by MT lattice, which have not been considered before, impose a strict directionality in conformational changes, causing different behaviors at the plus and the minus ends. Coexistence of multiple tubulin conformations in the MT lattice creates interfacial mechanical strains that spread over the entire MT and couple conformational changes over a wide region. This long-range coupling, which has not been considered before, makes conformational changes collective and leads to counter-intuitive emergent phenomena of catastrophe, rescue and pause.

Simulating all the phenomena of dynamic instability is technically challenging. We developed a novel strategy to properly integrate the mechanical strains with the kinetics of tubulin conformational changes, so that detailed balance is maintained. We also devised novel numerical strategies (details in the Supplemental Information (SI)) to drastically reduce the computational cost of kinetic simulations, allowing proper sampling of rare events with very slow time scales (e.g. catastrophe) in the context of a sea of frequent events (e.g., dimer association during growth), which makes the conventional Gillespie algorithm formidably expensive [34, 35].

Enabled by these novel concepts and technical innovations, our model quantitatively reproduces all phenomena of dynamic instability in simulation, passing the stringent test that no mechanistic elements are missing and no logical inconsistencies are present. The great variability in results across different groups suggests that quantitative behaviors of dynamic instability are very sensitive to experimental conditions [8, 11, 16]. Thus a rigorous test of a model needs to reproduce results from a consistent experimental setup. The classic experiment by Walker et al [8] remains the first and only comprehensive quantification of all phenomena of dynamic instability, including detailed minus-end behaviors, within a consistent experimental setup. Thus we choose ref. [8] as our simulation target. Simulations based on our model reproduced all the results in ref. [8], including pausing and minus-end dynamic instability, which have not been successfully modeled in previous studies. Below we first derive the key elements of our model from steric analyses of existing structures of tubulin, then detail the mechanisms for individual phenomena determined from simulations.

### II. Overview of known tubulin structures

Three polymer structures are observed in different phases of dynamic instability: 1) tube structure of stable MTs; 2) slightly curved sheet at growing plus-ends; and 3) highly curved PFs (“rams horn”) at depolymerizing ends. Each polymer structure consists of dimers in a conformation that complies with the polymer geometry. Dimers in the tube adopt a straight conformation (S form): axes of the monomers align perfectly and the intra-dimer interface is tight [18], and dimers in the rams horn adopt a curved conformation (C form): axes of the monomers form a 12° angle and the interface is looser, with reduced contact area and larger cavities (Fig. 1a) [19]. Structure of dimers in sheet derives from cryo-EM [11, 21] structure of cold-stable ribbons of GMPCPP-tubulins and has a 5° angle between two monomers, implying it is an intermediate between S and C. We call this conformation bent (B form). Muller-Reichert et al measured radius of curvature for short PFs of GDP and GMPCPP tubulins respectively [20]. The radius for GDP-tubulins suggests an angle of ~12° between two monomers, while the radius for GMPCPP-tubulins indicates an angle of ~6°±1.5°. Thus we assume that a GTP-tubulin in a PF without lateral bonds adopts B form. An intriguing finding in the ribbon structure is different lateral bonding on the two sides of a PF [21]. One is the same as those in tube (tube-like); the other is weaker (sheet-like). Converting sheet-like into tube-like lateral bonds, which occurs during sheet closure into tube, requires rotating a dimer around its longitudinal axis, feasible only for S form.

**Figure 1:**
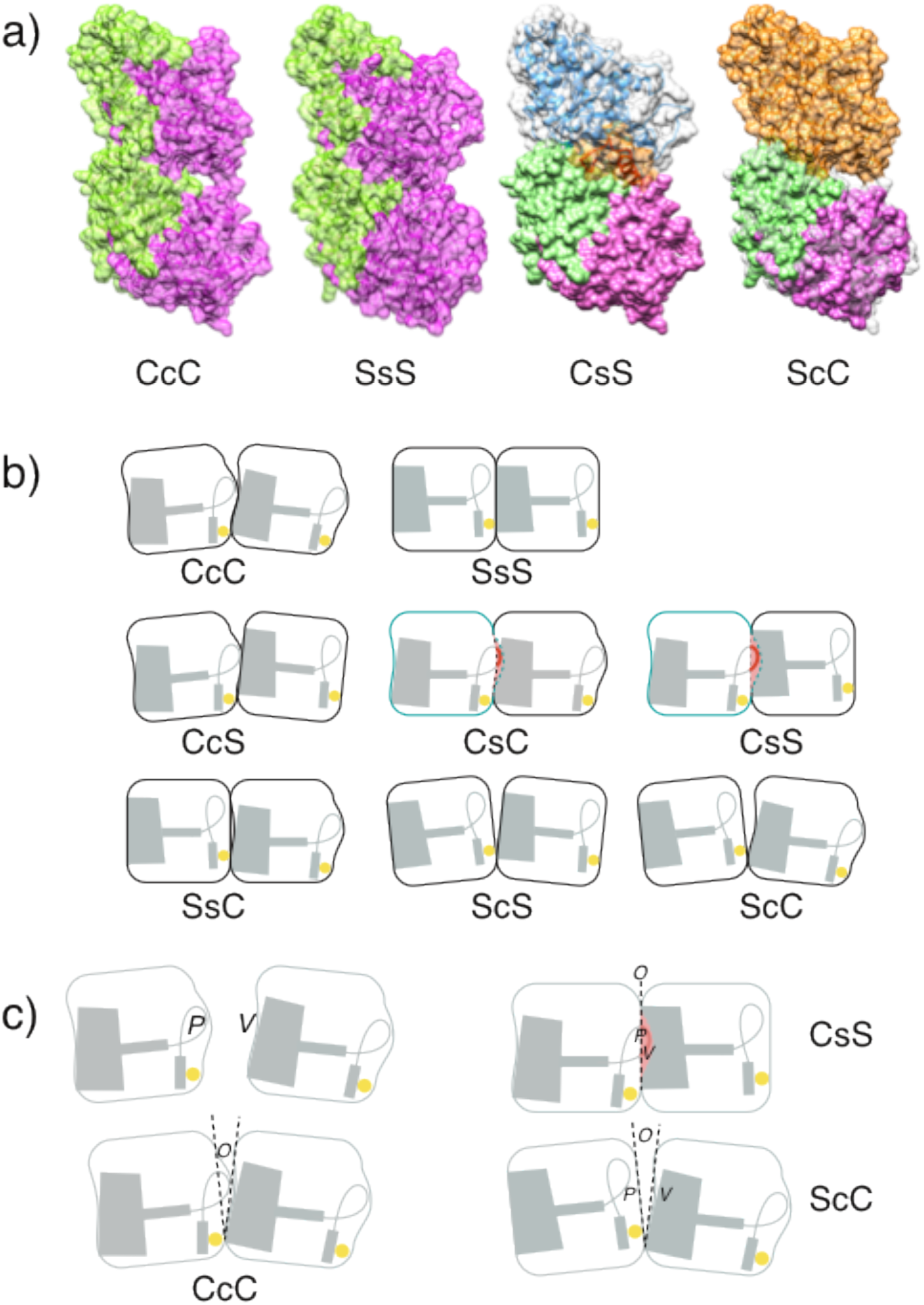
Structures and schematics of different dimer conformations. We use upper case for monomer conformations (i.e. S, B, C) and lower case for interface (i.e. s, b, c) conformations: SsS denotes a dimer or an intra-dimer interface in S form, where the left side is the minus-end side. Both sides of an inter-dimer interface are hyphenated (e.g. S-s-S). Since hydrolysis only modifies the nucleotide of β-monomer, we use a superscript for the nucleotide state of the β-monomer when necessary: T for GTP and D for GDP, so SsS^D^ represents a GDP-bound dimer. We use “e” for sheet-like and “t” for tube-like lateral bond, so BeB means two monomers that share a sheet-like lateral bond. (a) In the SsS (PDB: 1JFF) and CcC (PDB: 1SA0) structures in surface representation, N- and C-terminal domains are Green, the ID is Magenta. To construct CsS structure, the *β*-monomer in SsS structure is replaced by a *β*-monomer in C conformation (1SA0), aligned to it by N- and C-terminal domains. Some residues of the helix and loop in Red (their surface highlighted in Orange) clash with the *α*-monomer, as first discussed in ref. [19]. To construct ScC structure, the *α*-monomer (light Grey semi-transparent) in CcC structure is replaced by a *α*-monomer (solid Green and Magenta) in S conformation (1JFF) aligned to it by N- and C-terminal domains. The *β*-monomer is in Orange. A surface region continues from Orange to Gray represents contacts present in CcC but lost in ScC. (b) The basic notion of the schematics follows that of ref. [19]. The Red lines indicate steric strain or clash in the relevant dimer conformations. (c) Schematic representation of P, V and O.

### III. Steric analyses of tubulin structures and model assumptions

Structures above suggest a three-level hierarchy: 1) monomer and interface (longitudinal and lateral), 2) dimer, and 3) polymer structures. Stable tubulin structures must be consistent across this hierarchy and the nucleotide state decides the stable polymer structures: tube and rams horn are preferred by GTP and GDP-tubulins, respectively. Since different polymer structures dominate different phases of dynamic instability, numerous tubulin conformational transitions among B, S and C forms must occur, and the transitions need to proceed one monomer or interface at a time, each a separate first order chemical reaction. Thus a MT must go through a sequence of structural intermediates, each a combination of monomer and interface structures from both the reactant and the product polymer structures. The energies of these structural intermediates dictate the kinetics of dynamic instability by determining kinetic pathways of tubulin conformational changes.

According to the reasonings above, dynamic instability is dominated by tubulin conformational transitions that drive toward a stable polymer structure decided by nucleotide state. In the growing phase, a new dimer binding from solution forms a longitudinal bond with the tip of an existing MT, resembling a dimer in a GMPCPP-tubulin PF [20], and adopts the B form. It then forms lateral bonds with neighboring dimers at the tip, integrating into a sheet [11, 21]. This sheet closes into a tube after the dimers undergo B→S transition and their lateral bonds convert [11]. After dimers adopt the S form, GTP hydrolysis starts [18], enabling the S→C transitions required for catastrophe. Once C forms have swept the tip row, they propagate along the PFs, initiating rapid shortening. Lateral bonds then break because they are geometrically incompatible with the C form. As the C form advances, curved PFs cleave stochastically, exposing dimers in S form, which can bind new dimers and initiate rescue. We call this the B→S→C (or BSC) model from the order of these transitions.

The key to dynamic instability thus lies in tubulin conformational changes. Due to the consistency across the hierarchy of tubulin structures, all of the critical information is encoded in structures of the monomers and interfaces. Qualitative rules governing the energies of the structural intermediates can be deduced from a steric analysis of these structures.

#### a. Key features of monomer and interface structures

The structure of a monomer consists of a scaffold and a sliding part [19], where the scaffold contains N- and C-terminal domains and the sliding part is the intermediate domain (ID) [18, 19]. The positioning of the ID relative to the scaffold distinguishes monomer conformations. In the S form, the ID is embedded in the scaffold, creating flat surfaces on both sides of a monomer. This allows the plus-end surface of an α-monomer and the minus-end surface of a β-monomer to complement each other in a tight straight (S) interface [18]. We define the conformation of a longitudinal interface by the angle (*γ*) between two monomers: the S interface has *γ* = 0°. In the C form, the ID slides towards plus-end side, creating a protrusion (P) in the plus-end surface and a concavity (V) in the minus-end surface (Fig. 1c), where P is more pronounced than V [19]. If the interface between two monomers in C form adopts an S form, V on the plus-end side will not accommodate P on the minus-end side and steric strain ensues (Fig. 1c). Scaffolds of the two monomers have to rotate to open up the spacing, so V can accommodate P. The resulting interface is C with *γ* = 12°.

#### b. Steric analysis of interface energies

The interaction energy between two monomers forming an interface is the interface energy. It is determined by conformations of the interface and its monomers, as well as the nucleotide state. These effects can be deduced from geometric and steric considerations. The size of P and V decides how monomers on the minus- and plus-end side contribute to interface energy, respectively, while how the interface state contributes is determined by the extent of its opening (O) (Fig. 1c). Sizes of structural features P, V and O rank as f_C_>f_B_>f_S_ [18, 19], where f = P, V, O and the subscript (C, B, or S) indicates the form and the rank of the feature size. For optimal interactions at an interface, the *V* + *O* pocket should accommodate and complement P: *V*_*c*_ + *O*_*c*_ optimally complements *P*_*c*_, and *V*_*s*_ + *O*_*s*_ optimally complements *P*_*s*_. If *V* + *O* cannot accommodate P, a steric strain or clash occurs (Fig. 1). In a CsS (notations in Fig. 1) configuration, monomers clash because both *O*_*s*_ and *V*_*s*_ are too small, as shown in ref. [19]. On the other hand, if *V* + *O* is larger than P, the contact between monomers is loose and the interaction is weak. Both steric strain and loose contacts raise energy. For simplicity, we do not distinguish intra- and inter-dimer interfaces, thus SsS and S-s-S interfaces are treated identically.

#### c. Steric rules for interface energy

Based on the steric analyses, we can derive simple rules on energies of different interface configurations. The size of *P*_*a*_, *V*_*a*_ and *O*_*a*_ (a = S, B, C) are ranked as: C > B > S. Optimal interface configurations occur when P, V, O are of the same rank (e.g. all are C). If P has a higher rank than either V or O, the structure has steric strain. If P has a higher rank than both V and O, the structure has steric clashes, and is energetically forbidden from appearing in any kinetic pathway of tubulin conformational changes. If either V or O is of higher rank than P, then the monomers interact weakly. If both V and O are of higher rank, the interaction is weaker.

#### d. Impact of nucleotide: inter-dimer interface energy

At an inter-dimer interface, interactions of the nucleotide with the β- and α-monomers contribute to monomer and interface energies respectively, because it binds to the former and interacts with the latter across the interface A new structure by Zhang et al has shown that hydrolysis leads to compaction of the inter-dimer longitudinal interface and conformational changes in α-tubulins [14, 36], indicating that the nucleotide mainly impacts structures of the interface and the α-monomer. Thus we assume the nucleotide alters the interface energy.

### Condition for GTP hydrolysis

GDP-tubulins are found in both MTs and Zn induced flat sheets [18], confirming that the S-s-S interface is sufficient for hydrolysis. Because enzymatic catalysis requires precise positioning of relevant residues and therefore specific interface conformations, we restrict hydrolysis to S^T^-s-S interfaces.

### Mechanical energy in the system

Since a MT is a polymer, deviations from its equilibrium structure can have long-range effects, altering the reaction kinetics of monomers on many different rows. Thus we divide the MT energy into chemical and mechanical components. The chemical component accounts for changes in local chemical states (i.e. conformations, bonding, nucleotide state) of monomers and interfaces, while the mechanical component accounts for long-range effects that conformational changes have on global polymer structure. The two energy components form a feedback loop, with the global chemical state of the polymer dictating the mechanical energy and the mechanical energy affecting the changes in local chemical states. For example, when a monomer in sheet converts into S form, its equilibrium geometry no longer agrees with the surrounding monomers in B form. Its longitudinal and lateral bonds with its neighbors create a mechanical strain that is distributed over many monomer-monomer interactions, affecting their conformational changes. The net result is to favor conformational changes that reduce mechanical strain and disfavor those that increase it, with the extent determined by how much they reduce or increase it. We use harmonic terms for the mechanical energy (details in the SI).

## IV. Simulation Results

The steric analysis above provides sufficient information for pinning down kinetic pathways for all the tubulin conformational changes in dynamic instability. These pathways are optimized from exploring all possible options guided by the steric rules and considering the steric constraints of the MT lattice. Below we discuss in detail these pathways and how they dictate different phenomena of dynamic instability.

### 1. Plus-end growth and catastrophe

During growth and catastrophe, plus-ends display four key behaviors [7, 8, 11] (Fig. 6): 1) bi-phase behavior: a growing plus-end alternates between sheet and tube; 2) catastrophe; 3) pausing; and 4) a linear increase of growth rates with tubulin concentration. At the heart of these behaviors are B→S and S→C transitions.

### B→S transition pathway at plus-end: sequentially forward

When dimers bind to the tip of a plus-end, they adopt the B form and integrate into a sheet. The sheet then converts into tube via progressive B→S transitions. Since B is an intermediate between C and S, the differences between C and S indicate that the B→S transition of a monomer amounts to sliding the ID towards the minus-end side (Fig. 1). This directionality in the monomer transition imposes a strict directionality and ordering on the B→S transitions along a PF.

The B→S transition starts with a BbB dimer configuration (Fig. 2a). If the β-monomer transitions to an S form first, a BbS interface ensues, which has steric strain according to our steric rules. If the interface transitions next, a BsS configuration follows, which has steric clashes and is forbidden. Thus the B→S transition cannot start from the β-monomer and move sequentially towards the α–monomer. In contrast, if the α-monomer adopts S form first, an SbB interface results, which has loose contact but no steric strain, making it more stable than BbS. The interface transitions next, resulting in SsB, which has tighter contact than SbB and is more stable. Then the β-monomer changes, resulting in SsS, the optimal configuration for rank S. Finally, the two SeS lateral bonds of the dimer convert to StS, and the energy of the final state (SsS) decreases below that of the initial state (BbB). This pathway has a low starting barrier and is energetically downhill in subsequent steps (Fig. 2a). It has two features: 1) a preferred direction of propagation from minus-end towards plus-end; and 2) structural elements undergo B→S transitions in a strictly sequential order. With this pathway, the B→S transition initiates from the tube-sheet boundary and propagates sequentially towards the plus-end to close sheet into tube––its directionality and ordering conform with that of plus-end growth.

**Figure 2:**
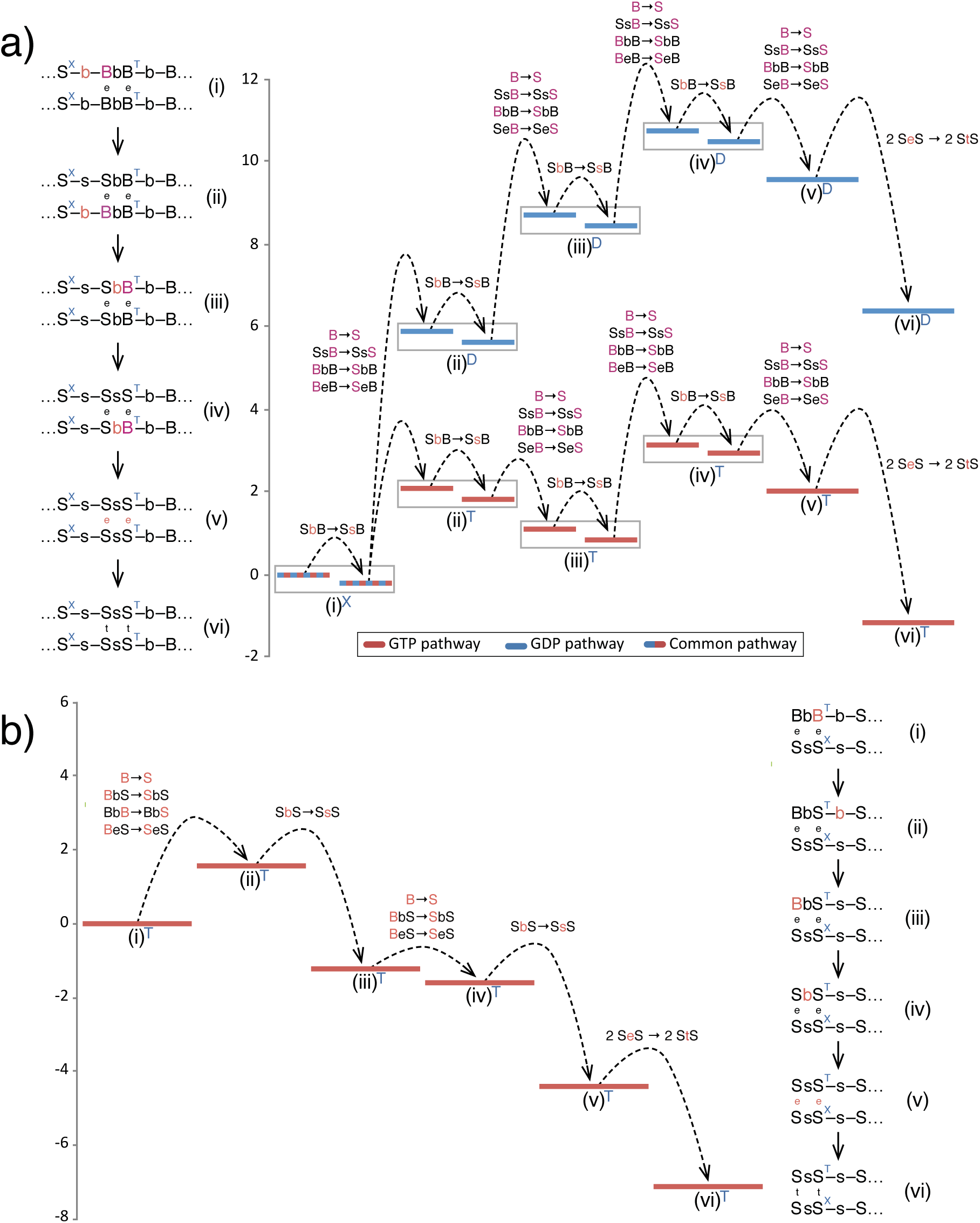
Optimal kinetic pathways and schematic energy diagrams for B → S transition. (a) B → S transition at plus-end with GTP and GDP at the tube-sheet boundary respectively. (b) B → S transition at minus-end.

### Growth and B→S transition: bi-phase from dual rates

Plus-end growth is biphase because there are two rates for B→S transition. Since the S^T^-s-S interface is more stable than the S^D^-s-S interface (Fig. 2a), S^T^-s-B→S^T^-s-S is faster than sheet extension at high tubulin concentration, while S^D^-s-B→S^D^-s-S is slower than sheet extension at low concentration, due to its higher energy cost. This feature enables sheet and tube phases to alternate at the plus-end with clear signatures for entrance and exit. To grow in sheet, the tube-sheet boundary needs to be in S^D^-s-B configuration, which traps dimers at the boundary in B form due to the slow S^D^-s-B→S^D^-s-S transition. Thus all the subsequent dimers are also trapped in B form because B →S transition must proceed sequentially. We call this S^D^-s-B configuration a “GDP-trap”. Once the S^D^-s-B→S^D^-s-S transition occurred after long enough waiting time, the subsequent B→S transitions proceed with the fast rate and overtakes the front of sheet, and the MT exits sheet to enter the tube phase.

### Tube to sheet: forming the GDP-trap

Once in tube phase, newly added dimers convert into S form before another dimer binds to it, keeping the tip growing in tube form. Meanwhile, a GDP-trap can form by chance. When a new dimer adds to a tip with a GTP-dimer in S form, the new dimer can begin fast B→S transition even before forming lateral bonds. When the inter-dimer interface between the new dimer and the original tip becomes S-s-S (Fig. 5a), GTP hydrolysis at this interface is enabled. After the hydrolysis, if this tip dimer fluctuates backward—an S→B transition—two scenarios can occur. In one case, lateral bonds form during S→B transition, preventing dissociation at the S^D^-b-B (Fig. 5a) interface and creating a GDP-trap. Alternatively, lateral bonds do not form and the tip dimer dissociates easily at the less stable S-b-B or S-s-B interface, exposing the original tip but now bound to GDP. When dimers bind to this tip and form lateral bonds, a GDP-trap is formed. Once GDP-trap is formed, growth proceeds as a sheet and the MT leaves tube phase.

While the duration of tube phase is determined by the time to form GDP-traps, the duration of sheet phase is determined by two factors: 1) the reaction time of S^D^-s-B→S^D^-s-S transition; and 2) the relative rates of S^T^-s-B→S^T^-s-S transition and sheet growth. While waiting for S^D^-s-B→S^D^-s-S transition, the plus-end grows by adding dimers in sheet form, so both MT and sheet elongate (Fig. 5b). Once the S^D^-s-B→S^D^-s-S transition completes, the B→S transition becomes fast and the front of S form chases the sheet tip. Thus the MT elongates but its sheet shortens (Fig. 5b, S5).

### Interface conformations that determine plus-end growth rate

Since conformational changes are essential to growth, growth rates are controlled by interface conformations. Because the MT lattice is two-dimensional, dimers incorporate into (2D-association) and dissociate from (2D-dissociation) a MT by multi-step composite processes. In 2D-association, a dimer binds to a MT tip at a rate proportional to tubulin concentration. It either dissociates again in an unproductive binding event or forms lateral bonds and incorporates into the lattice. The rate for 2D-association is the net rate for incorporating dimers into the lattice and dictates the slope of the concentration dependence of growth rates. In 2D-dissociation, dimers in the MT lattice, which are independent of tubulin concentration, need to break both longitudinal and lateral bonds. This process dictates the y-intercept of the concentration dependence of growth rates. In tube phase (details in the SI), S-b-B, StS, and SeS interfaces determine 2D-association, whereas StS and S-s-S control 2D-dissociation. In sheet phase, B-b-B, BeB, and BtB determine both 2D-association and 2D-dissociation. Figure 6a shows that our simulations reproduced the plus end growth rates in ref. [8].

### Catastrophe: a tug of war

Since all dimers in sheet are GTP-bound, catastrophe only occurs in tube phase, when a tip dimer is under constant competition between dimer addition and S→C transition. When dimer addition prevails, the MT grows in tube or sheet. When S→C prevails, catastrophe can occur.

#### a. S→C transition pathway at plus end: sequentially backward

The directionality of the S→C transition of a dimer dictates S→C transition along a protofilament. Unlike B→S, the S→C transition of a dimer cannot start from the α-monomer because it would create a CsS interface (Fig. 1), which has steric clashes and is energetically forbidden [19]. Instead, the optimal pathway starts from the β-monomer at the tip of the plus-end and propagates sequentially backward towards minus-end (Fig. 3), in the same direction as shortening at the plus-end.

**Figure 3:**
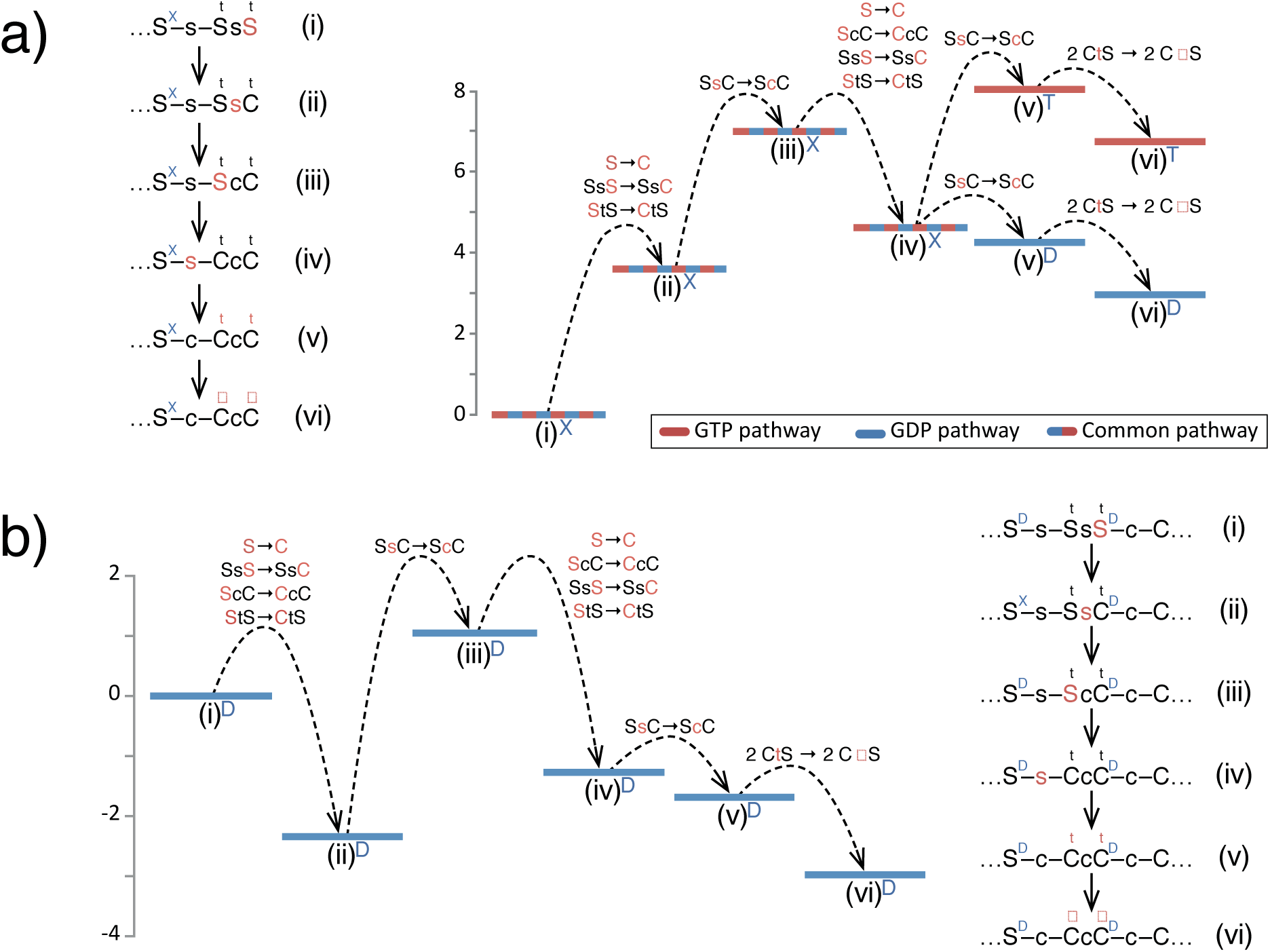
Optimal kinetic pathways and schematic energy diagrams for S → C transitions at plus-end. (a) S → C initiation with GTP and GDP behind the tip dimer respectively. (b) S → C propagation.

#### b. S→C initiation: dictated from behind

We call the S→C transition of dimers at the tip S→C initiation. Since a nucleotide only affects interface energy, S→C initiation is not affected by the nucleotide bound to the β-monomer at the tip, but is instead controlled by the nucleotide at the inter-dimer interface behind the tip dimer. When this interface is GDP-bound, the second step (SsC→ScC) (Fig. 3a) marks the highest energy on its S→C pathway, and steps afterwards are energetically downhill. In contrast, when this interface is GTP-bound, the step S^T^-s-C→S^T^-c-C (Fig. 3a) causes a steep increase in energy, making S→C initiation impractical.

#### c. S→C initiation and dimer addition: mutual exclusion

After a dimer at the tip adopts C form, no new dimer can bind to it because they cannot form a stable interface. The only options are C-b-B and C-c-B, both are unstable: C-b-B suffers steric clashes (see CsS in Fig. 1b) and C-c-B has steric strain. Therefore, S→C initiation excludes further dimer addition. Similarly, when a dimer at the tip binds a new dimer, S→C transition can no longer proceeds at the original tip. In this way, dimer addition excludes S→C initiation. Therefore, dimer addition and S→C initiation directly compete and are mutually exclusive.

After a tip dimer converts into C form, its lateral bonds will break because CtS interface has high energy, causing two effects. 1) The dimer in C form is now linked to the lattice by a single longitudinal bond and can easily dissociate and expose the next S dimer in line, which can bind a new dimer and resume growth (a microscopic rescue event). 2) Without lateral bonds, neighbors of the C dimer convert to C form more easily, causing S→C initiation to laterally propagate through the tip row. When the entire tip row converts into C form, catastrophe occurs.

Catastrophe is a direct competition between S→C initiation and adding new dimers. While the former does not depend on the tubulin concentration, the latter does. At higher concentrations where growth is fast, S→C initiation succeeds less often, making catastrophe frequency decrease with increasing tubulin concentration (Fig. 6b). Our simulations reproduced catastrophe frequencies in ref. [8] (Fig. 6b).

### Pausing: stuck at the GDP-tip

A peculiar puzzle is pausing, where a plus-end neither grows nor shortens [8]. Our model poses a natural mechanism for this behavior: pauses are transitional states between tube and sheet. Pauses happen during growth in tube form with a blunt tip geometry, when enough tip dimers become GDP-bound that growth stalls. We call a tip dimer that is GDP-bound and in S form a “GDP-tip.” Dimers added to GDP-tips have an S^D^-b-B interface (i.e. GDP-trap), and cannot quickly undergo the S-b-B → S-s-B → S-s-S transitions. Neither can they form lateral bonds easily due to mechanical penalty: sheet-like lateral bonds force the dimers into an orientation incompatible with that of the dimers on the tube side. This mismatch causes the longitudinal bonds between the tube and sheet dimers to twist, increasing mechanical energy and thus hindering lateral bond formation. The difficulty in forming lateral bonds increases with the number of GDP-tips. Consequently, although the rate of dimer association is unchanged, the rate of dissociation increases due to the unstable S-b-B interface, which causes plus-end growth to stall, leaving the MT paused. Eventually dimer addition succeeds when a pair of adjacent dimers associate and form a lateral bond before either dissociates––they are stabilized and initiate growth in sheet phase. In essence, a pause occurs when a MT is trapped with enough GDP-tip states and cannot grow. While paused, the high dissociation at S-b-B also reduces competition of dimmer addition with S→C initiation—an important mechanism for augmenting catastrophe. However, a sheet can often initiates from a few GDP-tips with no observable pause. Thus pausing is a rare event (Fig. 5b).

#### 2. Plus-end shortening and rescue

The most distinctive features during shortening are [7, 8]: 1) the shortening rate is much faster (up to 30 times) than the growth rate; 2) the frequency of rescue is much higher than catastrophe despite the faster shortening rate. These arise from the difference between S→C propagation (the S→C transition of dimers in the middle of PFs) and S→C initiation.

#### S→C propagation: allostery expedites shortening

The main process of shortening is S→C propagation. Its first step is almost the same as S→C initiation: the β-monomer converts into C conformation. But this step causes two concerted changes (Fig. 3b): 1) SsS→SsC, and 2) S-c-C→C-c-C, whereas S→C initiation has only the first one. The first change is energetically uphill, while the second one is downhill, thus energy gained from the second change compensates the cost of the first one, making propagation easier than initiation. This effect resembles allostery in T→R transition of hemoglobin [37]—the cost of a monomer structural change is compensated by energy gained from the resulting change in the interface between monomers. This allosteric effect makes S→C propagation much faster than initiation and explains why catastrophe is a rare event while shortening is a fast process. Figure 6c shows that our simulations reproduced shortening rates in ref. [8].

### Rescue: the inverse of catastrophe

The final steps of S→C transition of a dimer create S-s-C or S-c-C interface with the next dimer in the PF. Both interfaces, especially S-c-C, are metastable because they have loose interactions. Longitudinal bonds break easily at these interfaces, and as rapid shortening proceeds, there is a direct competition between S→C propagation and S-c-C bond breaking. If the S-c-C bond breaks, the curved rams’ horn of that PF is cleaved and dissociates, leaving the tip as a GDP-bound dimer in S form (i.e. a GDP-tip), which mirrors the growing tip in tube phase. The exposed dimer can bind a new dimer or undergo S→C initiation. Until it undergoes S→C initiation, rapid shortening effectively stops. If several adjacent rams’ horns cleave, then the new dimers can start to associate, causing a rescue event. Since rescue occurs on GDP-tips, it often renews growth in sheet or occasionally manifests as pausing during shortening (Fig. 5b) [8].

In rescue, growth competes against S→C initiation rather than propagation. Because initiation is much slower than propagation (Fig. 4c), the chance for resuming growth is high. This explains the puzzlingly high frequency of rescue when shortening is up to thirty times faster than growth (Figs. 6c, d). Rescue is determined by two factors: 1) the rate that a depolymerizing PF cleaves, and 2) the chance that a newly exposed S dimer binds a new dimer. The first determines how often a PF qualifies for rescue; the second determines how likely a qualified PF rescues and is the reverse of what determines catastrophe. Figure 6d shows that our simulations reproduced rescue results in ref. [8].

**Figure 4:**
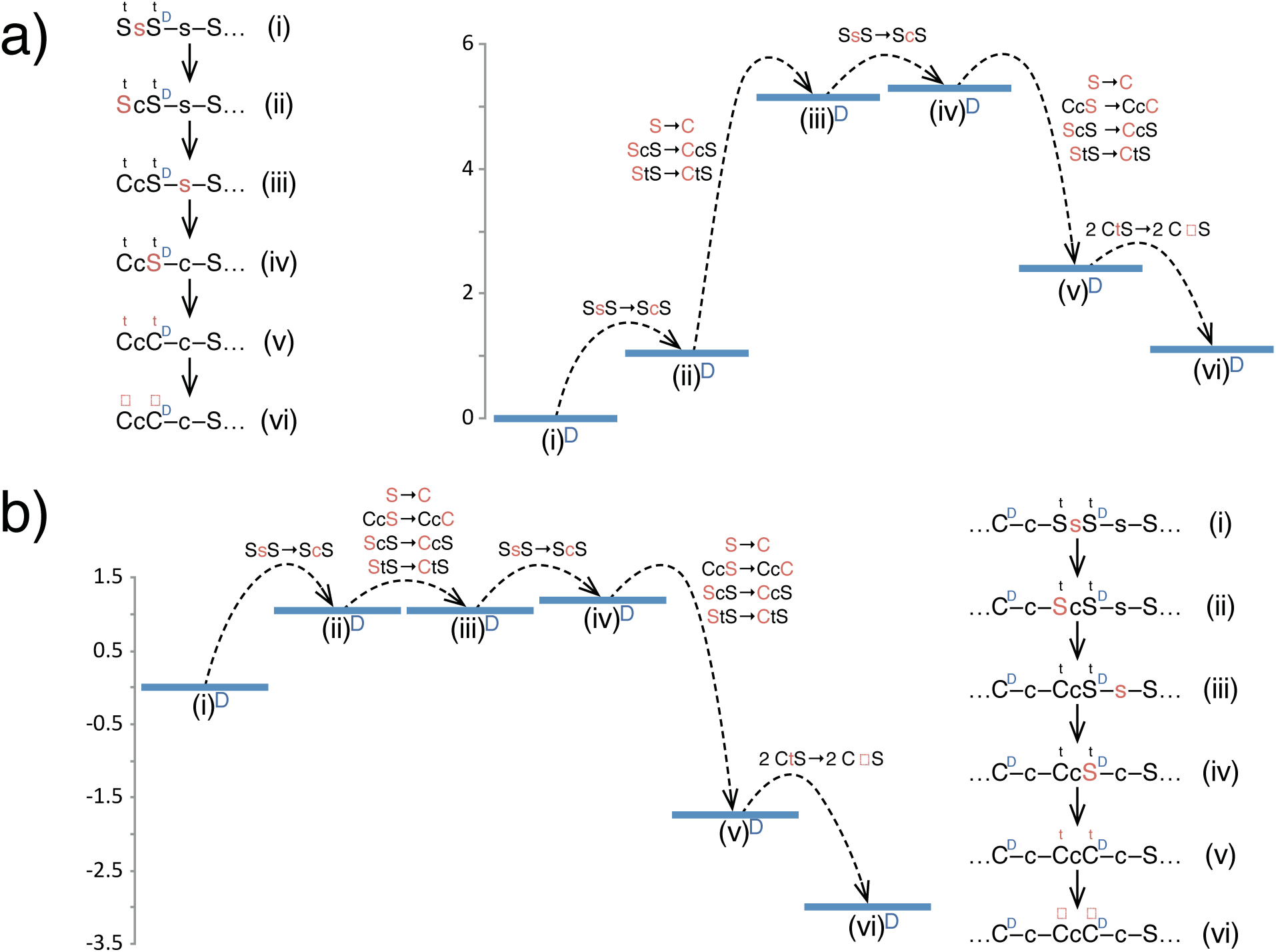
Optimal kinetic pathways and schematic energy diagrams for S → C transitions at minus-end. (a) S → C initiation. (d) S → C propagation.

**Figure 5:**
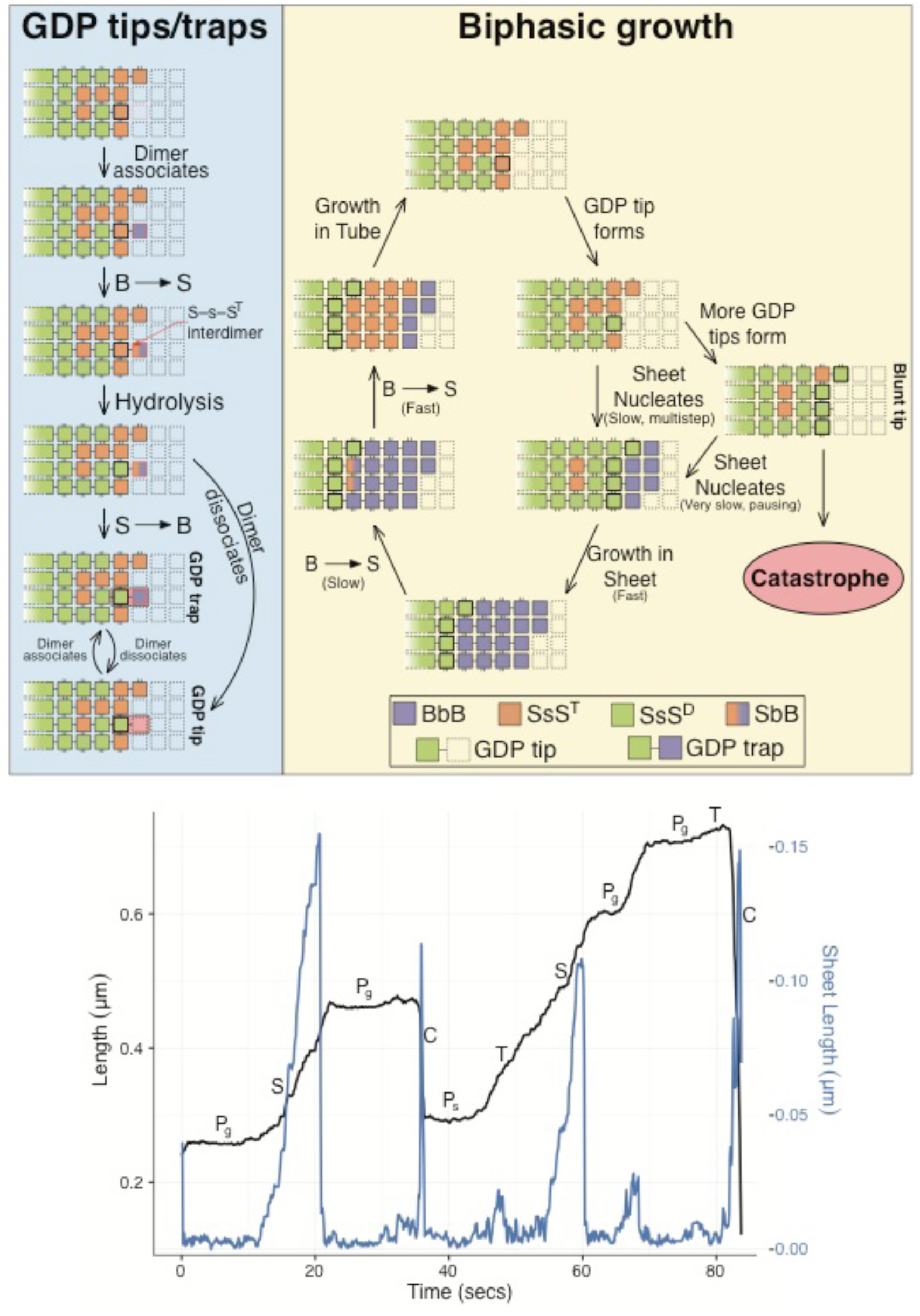
Illustrations of dynamic instability at plus-end. (a) A schematic for the mechanism of bi-phase growth at plus-end. (b) An example trajectory from plus-end simulation with different phases marked. S: growth in sheet; T: growth in tube; P_g_: pausing during growth; P_s_: pausing during rapid shortening; C: catastrophe.

### Reciprocity between catastrophe and rescue

Catastrophe and rescue are determined by the same competition between dimer addition and S→C initiation, but at growing and shortening tips, respectively. They share the same underlying processes, but with opposite definitions for success. If S→C initiation wins, the result is catastrophe; if dimer addition wins, the result is rescue. This reciprocity manifests in two aspects. 1) The extra cost for S→C initiation, compared to propagation, makes catastrophe harder and rescue easier––recue is eighty times more frequent than catastrophe. 2) Dimer addition increases with tubulin concentration, thus it increases rescue but decreases catastrophe.

### 3. Minus-end growth and catastrophe

Growth at the minus-end is much slower than at the plus-end and catastrophe is also less frequent [8]. What could cause these discrepancies given the same molecular interfaces are at both ends? The answer lies in the directionality of B→S and S→C transitions at the minus-end.

#### B→S transition pathway at minus-end: a back-and-forth sequence

Similar to the plus-end, dimers bind to the minus-end in B form and then convert to S form. But the B→S transition cannot follow the sequential pathway of the plus-end because it proceeds in the opposite direction of minus-end growth. Instead, in the optimal minus-end pathway (Fig. 2b) conformational changes occur in a back-and-forth pattern that conforms to the direction of growth while avoiding steric clashes and minimizing energy barriers. With this pathway, B→S transition propagates from the plus-end towards the minus-end, a direction opposite to the plus-end pathway.

#### B→S transition and minus-end growth: one rate, one phase

Contrary to plus-end, the nucleotide at the interface between a new dimer and the tip of a minus-end is always GTP, because it is brought in by the new dimer and cannot hydrolyze before the new dimer completes a B→S transition. Therefore, unlike the plus-end, the minus-end has only one rate for B→S and cannot show bi-phase behavior. Instead, it grows only as a tube, requiring B→S to be faster than dimer addition at all concentrations.

#### Killing two birds with one stone: the B-b-S binding energy slows growth and expedites B→S

The initial interface between a new dimer and minus tip is B-b-S, which has high energy due to steric strain. Therefore, the new dimer tends to dissociate before forming lateral bonds, decreasing minus-end growth. In addition, B-b-S marks the highest energy on the B→S transition pathway (Fig. 2b). This high cost is fully paid by the binding energy of the new dimer and in subsequent steps the B-b-S interface is retained, but never created *de novo*. Once the initial cost of the B-b-S interface is paid by binding, it does not need to be paid again, and subsequent steps in minus-end B→S are energetically downhill and rapid. This mechanism both slows growth and speeds B→S transition, making minus-end grow in tube.

As discussed above, the interfaces responsible for the 2D-association at the minus-end are B-b-S, StS and SeS. The interfaces governing minus-end 2D-dissociation are StS and S-s-S. Figure 6a shows that simulations captured the minus-end growth rates in ref. [8].

**Figure 6:**
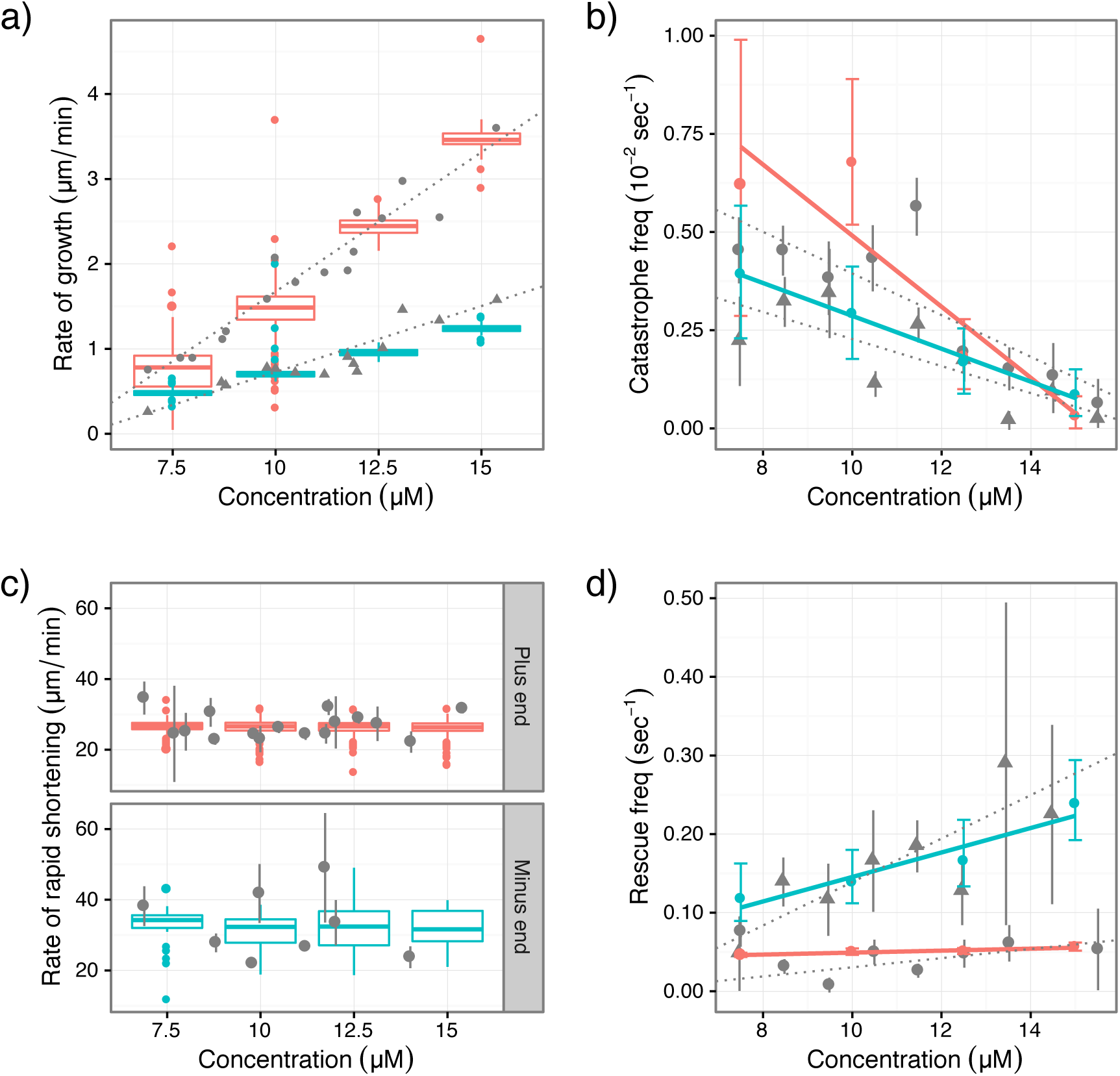
Comparison between results of simulations and experiments in ref. [8]. The experimental data are obtained by digitization of figures in ref. [8]. Experimental data are in gray; triangles denote plus-end data and dots denote minus-end data; vertical lines are error bars and dotted lines are linear fitting from ref. [8]. For simulation results, plus-end is red and minus-end is cyan. Growth and shortening rates are presented as standard box plots; frequencies for catastrophe and rescue are presented as error bars. Simulations are carried out at tubulin concentration of 7.5, 10.0, 12.5 and 15.0 µM. Linear regression (solid lines) was conducted on catastrophe and rescue frequencies. (a) growth rates; (b) catastrophe frequencies; (c) shortening rates; (d) rescue frequencies.

### Low catastrophe from a high cost of S→C initiation

Since minus-end grows in tube, dimer addition and S→C initiation are in competition at all times. The S→C pathway of the plus-end is disallowed at the minus-end because it first creates a C-s-S interface that is forbidden due to steric clashes (Fig. 1). Consequently, S→C transition at minus-end adopts a back-and-forth pathway (Fig. 4a). The C-c-S interface of this pathway is a high-energy state due to steric strain, making S→C initiation more difficult than at the plus-end. Although dimer addition at minus-end is slow, it is still faster than S→C initiation at the minus-end, leading to a lower catastrophe frequency than at the plus-end. Figure 6b shows that our simulations can reproduce catastrophe results in ref. [8].

### 4. Minus-end shortening and rescue

Contrary to the growing phase, minus-end has faster shortening and higher rescue than plus-end [8]. This puzzle is caused by the features of S→C propagation.

## Killing three birds with one stone: front-loading S→C creates fast shortening, low catastrophe and high rescue

The main event of minus-end shortening is S→C propagation towards plus-end. The highest barrier on this pathway (Fig. 4b) is ScS→CcS, which has a high cost due to the instability of CcS configuration. However, this cost is incurred only on a tip dimer, not on a dimer in the middle of a PF (Fig. 4b). During the minus-end S→C propagation, each step of the pathway maintains a C-c-S or CcS interaction by forming and annihilating C-c-S and CcS configurations simultaneously. Consequently, there are lower energy barriers for minus-end S→C propagation than for plus-end propagation, leading to faster shortening at the minus-end.

After a PF cleaves during shortening, the higher cost of S→C initiation at the minus-end makes dimer addition more likely to prevail than at the plus-end. Together they cause more frequent rescues at the minus-end. As shown in Fig. 6c and 6d, our simulations reproduced the minus-end shortening rates and rescue frequencies in ref. [8]. In summary, creating the high-energy CcS interface only once at the beginning of minus-end S→C transition has three effects: faster shortening, lower catastrophe, and higher rescue than at plus-end.

## Counter-intuitive experimental observations concerning the mechanism of plus-end catastrophe

Many ingenious experiments have been devoted to determine the size of GTP-cap at a growing plus-end and how the loss of GTP-cap leads to catastrophe, resulting in intriguing though counter-intuitive observations. These results are natural consequences of the mechanisms discussed in previous sections.

A major puzzle concerning the GTP-cap is the independence of its size on MT length: a small cap of up to three rows can prevent a long MT from catastrophe [12]. This finding contradicts the well-accepted idea that catastrophe results from GDP-tubulin-induced mechanical strain in MT lattice, which is counter-balanced by the restraining force of the GTP-cap. This mechanical strain should increase with MT length, thus a longer MT should require a longer GTP-cap. One remedy to this apparent dilemma is the implication of longer GTP-caps from EB1-based experiments [15], though this conclusion relies on the equivalence between EB1 binding region and GTP-cap, which requires further validation. Our model resolves this puzzle from a different perspective.

Although GDP-bound tubulins prefer the C form, it does not suggest that they cannot exist in the S form. Instead, it is more likely that, based on the standard picture of protein conformations and conformational dynamics [38], both the S and C forms are stable basins in the configuration space of tubulin regardless of the nucleotide state (Fig. 7a), but the S form is more stable in the GTP-bound state while the C form is more stable in the GDP-bound state. In either GTP- or GDP-bound state, a tubulin molecule can transit between S and C forms and this transition is a thermally activated process similar to a first-order chemical reaction. GTP hydrolysis flips the preferred direction of this transition, but does not make it occur and finish instantly.

**Figure 7:**
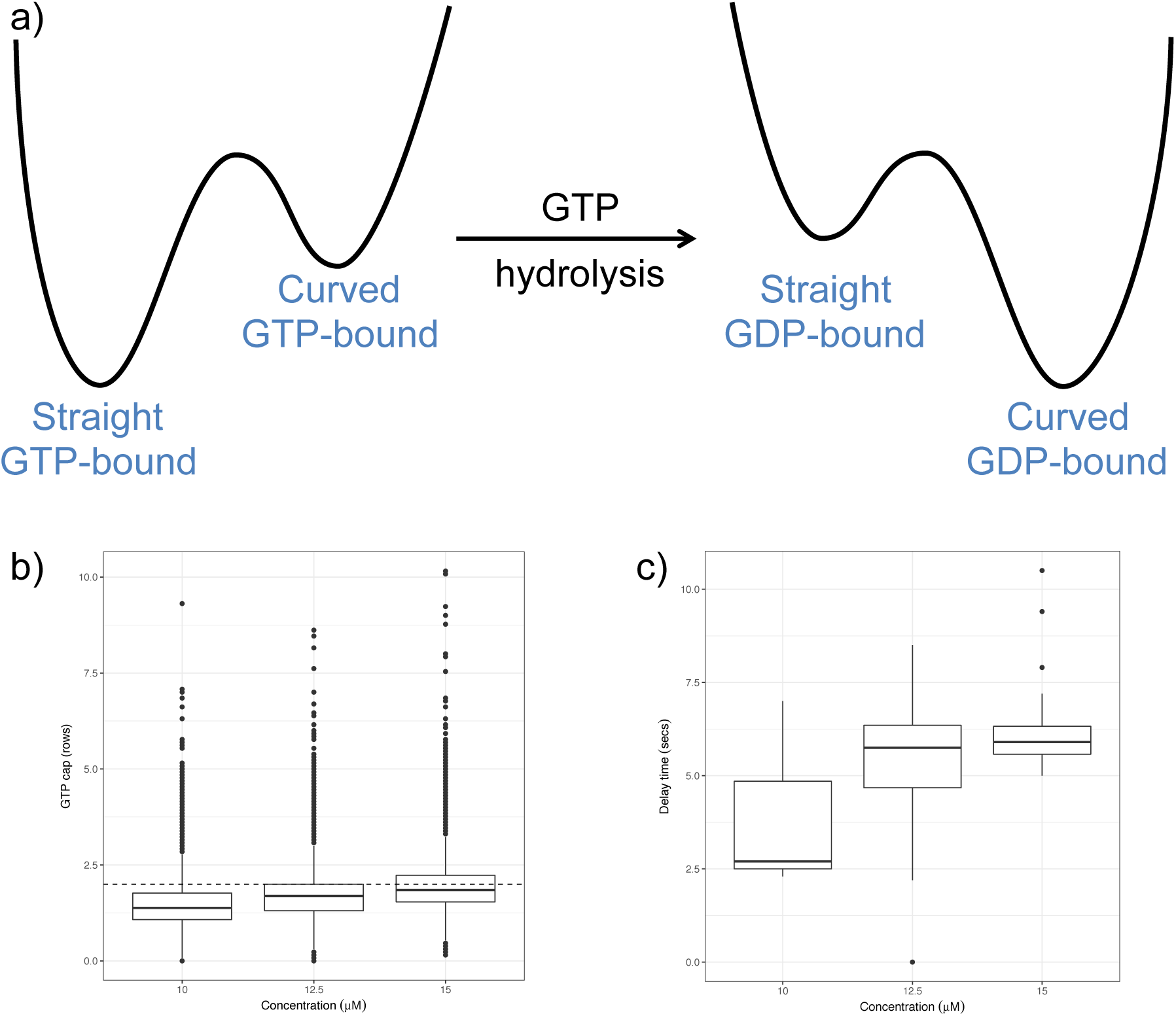
Results on the size of GTP-caps and simulations of the dilution experiment by Walker et al [41]. (a) A schematic diagram on the effect of GTP hydrolysis on tubulin conformation. In either GTP- or GDP-bound state, the S and C forms are two stable states separated by a transition barrier [38]. The stability of the S and C forms, indicated by the depth of the corresponding basin, changes with the nucleotide state. (b) Distributions of the size of GTP-caps during the tube phase of growth at different tubulin concentrations. The GTP-cap size is determined as the average number of rows of GTP-tubulins in S form at a growing plus-end. The dashed line marks the GTP-cap size of two rows. (c) Distributions of the delay time before a growing plus-end catastrophe upon sudden dilution with different pre-dilution tubulin concentrations. Dilution in simulations follows the procedure of Walker et at [41] by linearly decreasing tubulin concentration based on five points extract from the dilution curve in Fig. 4 of ref. [41].

Consequently, GDP-tubulins do not induce mechanical strains in the MT lattice. Since S→C transition must initiate from the MT tip and propagate sequentially towards the minus-end, a single tip dimer in S form holds an entire PF in S form. If the tip dimer stays in S form, all the dimers in the MT lattice stay in S form and do not incur the mechanical strain that drives catastrophe and depolymerization. This is consistent with recent findings by Driver et. al [39]. They used a laser trap to measure forces produced by depolymerizing MTs and found that shortening rate and mechanical strain are uncoupled to each other. The number of GTP-tubulins at growing plus-ends during tube phase in our simulations (Fig. 7b) all range between one and three rows (cases of GTP-caps longer than four rows account for only 0.25% of the data), an expected result since hydrolysis is a fast first-order reaction occurring at all S-s-S interfaces [2].

An important experimental observation supporting small GTP-cap is that growing MT plus-ends catastrophe within seconds after sudden dilution [40, 41]. This result is expected in our model given the short GTP-caps in our simulations during the tube phase, which is the dominant phase during growth because sheet phase accounts for less than 14% of the growth time. Indeed, in our simulations of the dilution experiment (details in the SI), plus-ends catastrophe very quickly (Fig. 7c), the same as observed by Walker et. al [41].

A final question is the kinetics of losing GTP-cap and committing catastrophe. The widespread idea casts catastrophe as a single-step process, but recent experiments proved it multi-step [16, 17], because the cumulative distribution of catastrophe time followed a Gamma-distribution (i.e. aging) instead of an exponential distribution expected from a single-step process. Recent modeling works provided different explanations into the nature of aging [6, 42, 43].

In our model, catastrophe is clearly multi-step. After the loss of both the structural cap (i.e. sheet closure into tube) and the GTP-cap (i.e. GTP hydrolysis at the longitudinal interface behind the tip row), catastrophe still requires that tip dimers in C form reach a critical number, which is analogous to the critical nucleus in condensation transition. This critical number is related to the shape parameter of the Gamma distribution and depends on experimental condition as catastrophe frequency does. Reaching this critical number requires multiple S→C initiations, making catastrophe multi-step. The rate of each S→C initiation is closely related to the rate parameter of the Gamma distribution and is not constant because it depends on the number of lateral bonds that need to break and the competition from dimer addition.

The cumulative distribution of catastrophe time in our simulations indeed follows a Gamma distribution (Fig. 8a), with a shape parameter 1.85 and a rate parameter 0.04 *s*^−1^. The time dependent catastrophe frequency (Fig. 8b) [16, 17, 44] also shows the typical increase with MT age. Because the MTs in Walker et. al [8], which is the target system of our simulations, behave very differently from the MTs in Gardner et al (e.g. MTs in ref. [16] do not rescue at all, have much lower catastrophe and much weaker concentration dependence of catastrophe frequency, and much slower growth rate), quantitative agreement between shape and rate parameters from our simulations and those from Gardner et al is not expected. On the other hand, our simulations captured the dependence of these parameters on tubulin concentration. The cumulative distribution of catastrophe time (Fig. 8c) shifts with increasing tubulin concentration in the same manner as shown in Fig. 3D of ref. [16]. The rate parameter decreases slightly with increasing tubulin concentration (Fig. 8d), an effect similar as that observed by Gardner et al [16]. This is a natural consequence of the mechanism of plus-end catastrophe in our model. The rate of dimer addition increases as tubulin concentration increases, making it more difficult for S–>C initiation to succeed. This slows down the effective rate of S–>C initiation and causes slight decrease of the rate parameter.

**Figure 8:**
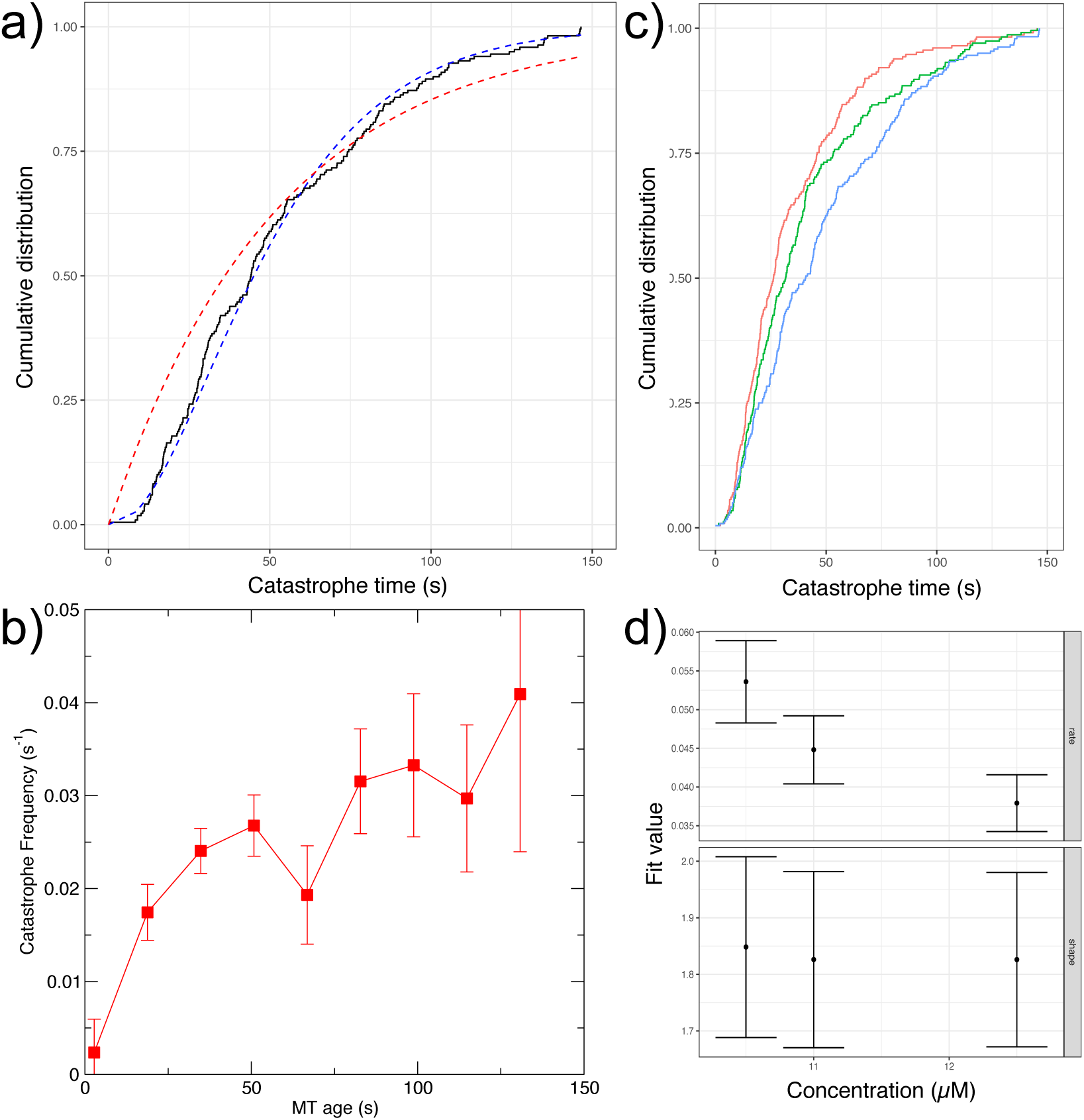
Simulation results on the aging phenomenon of plus-end catastrophe. (a) Cumulative distribution function (CDF) of catastrophe time at tubulin concentration of 12.5 µM. Black: CDF from simulations; Red dashed line: optimal exponential fit to the CDF; Blue dashed line: optimal Gamma distribution fit to the CDF (shape parameter: 1.85; rate parameter: 0.04 *s*^−1^). (b) Time dependent catastrophe frequency as a function of MT age, computed using Eq. (1) of ref. [16]. Error bars denote standard errors. (c) Comparison of CDFs of catastrophe time at tubulin concentrations of 10.5 µM (red), 11 µM (green) and 12.5 µM (blue). (d) Dependence of the rate (upper) and shape (lower) parameters of the best-fit Gamma distribution of CDFs of catastrophe time on tubulin concentration. All the fits were generated using the MASS::fitdistr function of R3.3.1.

## V. Discussion and conclusion

We have presented a mechano-chemical model that provides a coherent and unified mechanism for all the key phenomena of dynamic instability. In our model, dynamic instability is driven by conformational changes of tubulins: B→S transition dictates growth, and S→C transition dictates catastrophe, shortening and rescue. All the complex and counter-intuitive properties of these phenomena emerge aggregated effects of a key asymmetric feature of the structure of tubulin monomer: the intermediate domain of a monomer moves towards its minus-end side in B→S transition and its plus-end side in S→C transition. The directional arrangement of tubulin dimers in the MT lattice thus demands that B→S and S→C transitions at plus and minus ends must follow different kinetic pathways, which leads to different behaviors.

The B→S transition at plus-end must initiate from the tube-sheet boundary and propagate towards plus-end in a strictly sequential order to avoid steric clashes. Thus a dimer trapped in B form at this boundary holds all the subsequent dimers in a PF in B form (i.e. the sheet). Besides, B→S transition is much slower at a tube-sheet boundary bound with GDP (i.e. GDP-trap) instead of GTP, enabling the plus-end to alternate between sheet and tube phases during growth. It also causes pausing when dimers bind to MT tip bound with GDP (i.e. GDP-tip). In the B→S transition pathway at the minus-end, the state of the highest energy occurs when a new dimer binds to the minus-tip, whereas subsequent steps have low energy barrier. Thus a new dimer either dissociates or converts into S form quickly, leading to slow growth and fast B→S transition that keeps minus end in tube form all the time.

In contrast, the S→C transition at the plus-end must initiate from the MT tip and propagates towards the minus–end in a strictly sequential order; otherwise steric clashes ensue. Thus a tip dimer in S form can hold an entire PF straight. Besides, S→C transition is much easier for dimers in the middle of PFs than for dimers at the tip, because the former have allostery whereas the latter do not. This feature makes shortening fast and catastrophe rare, and enables rescue amid fast shortening. Compared to the plus-end, S→C transition at the minus-end has lower barriers for dimers in the middle of PFs but a higher barrier for dimers at the tip, leading to faster shortening, lower catastrophe, and higher rescue than the plus-end.

Some puzzling phenomena, such as small GTP-cap [12, 41, 45], fast catastrophe in response to sudden dilution [15, 40, 41], and aging [16, 17], are natural consequences of our model and can be understood coherently. Moreover, our model suggests an interesting dynamic mechanism for a most startling observation of MTs: the temperature-induced ribbon-to-tube conversion of GMPCPP-tubulins [21]. In our model, the mechanical energy poses a high transition barrier that makes the number of pathways over this barrier drop steeply with decreasing temperature (details in SI). The system is thus kinetically trapped in the metastable ribbon structure, and only at higher temperature are enough pathways available for the system to escape to its equilibrium tube structure.

Our model also provides a way to break through the bottleneck for further understanding the cellular MT network [1, 46, 47]. In this crucial cellular machinery, dynamic instability is the infrastructure and MT-associated proteins (MAPs) are fine-tuning controls. Plus-end tracking MAPs have attracted enormous attention [1, 4, 5, 46], and recently discovered minus-end MAPs have stimulated similar excitement [1, 48–50]. Despite ingenious *in vitro* experiments, the functions and mechanisms of MAPs remain elusive. Existing work defines MAP function by its impact on phenomenological processes. Our model suggests a different perspective: each phenomenological process consists of a series of elementary reactions that overlap with other phenomenological processes. The interactions of a specific MAP with a MT likely alter one or more of the elementary reactions, affecting all the processes that share these reactions. This explains why a single MAP often simultaneously affects opposite phenomenological processes (e.g. catastrophe and rescue) [51–53]. By identifying the underlying elementary reactions, our model provides a clean conceptual scaffolding to understand the mechanisms of MAPs. In addition, our model is the only one that explains dynamic instability at the minus-end, and should prove valuable in understanding the newly discovered minus-end MAPs that have only begun to be discovered.

## Methods

The simulation system is a MT of thirteen protofilaments represented as a two-dimensional lattice. Each non-empty lattice site is a monomer, a longitudinal interface, or a lateral interface. There are three types of chemical reactions: 1) conformational change of a monomer or interface; 2) forming and breaking a longitudinal or lateral bond; 3) GTP hydrolysis. Chemical reactions obey first order kinetics. The mechanical energy consists of harmonic terms to constrain geometric parameters to equilibrium values. There are five geometric coordinates, corresponding to stretching and bending of a longitudinal or lateral bond, and twisting around a longitudinal bond (details in SI). Chemical reactions are simulated using the Gillespie algorithm [34, 54] with a rejection step to incorporate the effects of the mechanical energy while preserving detailed balance. A chemical reaction selected by Gillespie algorithm is accepted with a ratio 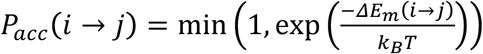, where *ΔE*_*m*_(*i* → *j*) is the change in mechanical energy induced by the tentative reaction through its perturbation to the polymer structure.

## References

1. Akhmanova, A. and M.O. Steinmetz, Control of microtubule organization and dynamics: two ends in the limelight. Nature Reviews Molecular Cell Biology, 2015. 16(12): p. 711–726.

2. Desai, A., Mitchison, T. J., Microtubule polymerization dynamcis. Ann. Rev. Cell Dev. Biol., 1997. 13: p. 83–117.

3. Kirschner, M. and T. Mitchison, Beyond self-assembly - from microtubules to morphogenesis. Cell, 1986. 45(3): p. 329–342.

4. Liu, J., et al., A mechanobiochemical mechanism for monooriented chromosome oscillation in mitosis. Proceedings of the National Academy of Sciences of the United States of America, 2007. 104(41): p. 16104–16109.

5. Liu, J. and J.N. Onuchic, A driving and coupling “Pac-Man” mechanism for chromosome poleward translocation in anaphase A. Proceedings of the National Academy of Sciences of the United States of America, 2006. 103(49): p. 18432–18437.

6. Bowne-Anderson, H., et al., Microtubule dynamic instability: A new model with coupled GTP hydrolysis and multistep catastrophe. Bioessays, 2013. 35(5): p. 452–461.

7. Mitchison, T.J. and M. Kirschner, Dynamic instability of microtubule growth. Nature, 1984. 312(5991): p. 237–242.

8. Walker, R.A., et al., Dynamic Instability of Individual Microtubules Analyzed by Video Light-Microscopy - Rate Constants and Transition Frequencies. Journal of Cell Biology, 1988. 107(4): p. 1437–1448.

9. Hill, T.L., Introductory Analysis of the Gtp-Cap Phase-Change Kinetics at the End of a Microtubule. Proceedings of the National Academy of Sciences of the United States of America-Biological Sciences, 1984. 81(21): p. 6728–6732.

10. Hill, T.L. and M.F. Carlier, Steady-State Theory of the Interference of Gtp Hydrolysis in the Mechanism of Microtubule Assembly. Proceedings of the National Academy of Sciences of the United States of America-Biological Sciences, 1983. 80(23): p. 7234–7238.

11. Chretien, D., S.D. Fuller, and E. Karsenti, Structure of Growing Microtubule Ends - 2-Dimensional Sheets Close into Tubes at Variable Rates. Journal of Cell Biology, 1995. 129(5): p. 1311–1328.

12. Drechsel, D.N. and M.W. Kirschner, The Minimum Gtp Cap Required to Stabilize Microtubules. Current Biology, 1994. 4(12): p. 1053–1061.

13. Caplow, M., R.L. Ruhlen, and J. Shanks, The Free-Energy for Hydrolysis of a Microtubule-Bound Nucleotide Triphosphate Is near Zero - All of the Free-Energy for Hydrolysis Is Stored in the Microtubule Lattice. Journal of Cell Biology, 1994. 127(3): p. 779–788.

14. Zhang, R., et al., Mechanistic Origin of Microtubule Dynamic Instability and Its Modulation by EB Proteins. Cell, 2015. 162(4): p. 849–859.

15. Duellberg, C., et al., The size of the EB cap determines instantaneous microtubule stability. Elife, 2016. 5.

16. Gardner, M.K., et al., Depolymerizing Kinesins Kip3 and MCAK Shape Cellular Microtubule Architecture by Differential Control of Catastrophe. Cell, 2011. 147(5): p. 1092–1103.

17. Odde, D.J., L. Cassimeris, and H.M. Buettner, Kinetics of Microtubule Catastrophe Assessed by Probabilistic Analysis. Biophysical Journal, 1995. 69(3): p. 796–802.

18. Nogales, E., S.G. Wolf, and K.H. Downing, Structure of the alpha beta tubulin dimer by electron crystallography. Nature, 1998. 391(6663): p. 199–203.

19. Ravelli, R.B.G., et al., Insight into tubulin regulation from a complex with colchicine and a stathmin-like domain. Nature, 2004. 428(6979): p. 198–202.

20. Muller-Reichert, T., et al., Structural changes at microtubule ends accompanying GTP hydrolysis: Information from a slowly hydrolyzable analogue of GTP, guanylyl (alpha,beta)methylenediphosphonate. Proceedings of the National Academy of Sciences of the United States of America, 1998. 95(7): p. 3661–3666.

21. Wang, H.W. and E. Nogales, Nucleotide-dependent bending flexibility of tubulin regulates microtubule assembly. Nature, 2005. 435(7044): p. 911–915.

22. Hill, T.L. and Y. Chen, Phase-Changes at the End of a Microtubule with a Gtp Cap. Proceedings of the National Academy of Sciences of the United States of America-Biological Sciences, 1984. 81(18): p. 5772–5776.

23. Bayley, P.M., M.J. Schilstra, and S.R. Martin, Microtubule Dynamic Instability - Numerical-Simulation of Microtubule Transition Properties Using a Lateral Cap Model. Journal of Cell Science, 1990. 95: p. 33–48.

24. Brun, L., et al., A theory of microtubule catastrophes and their regulation. Proceedings of the National Academy of Sciences of the United States of America, 2009. 106(50): p. 21173–21178.

25. Flyvbjerg, H., T.E. Holy, and S. Leibler, Stochastic Dynamics of Microtubules - a Model for Caps and Catastrophes. Physical Review Letters, 1994. 73(17): p. 2372–2375.

26. Li, X. and A.B. Kolomeisky, Theoretical Analysis of Microtubule Dynamics at All Times. Journal of Physical Chemistry B, 2014. 118(48): p. 13777–13784.

27. Margolin, G., et al., The mechanisms of microtubule catastrophe and rescue: implications from analysis of a dimer-scale computational model. Molecular Biology of the Cell, 2012. 23(4): p. 642–656.

28. Padinhateeri, R., A.B. Kolomeisky, and D. Lacoste, Random Hydrolysis Controls the Dynamic Instability of Microtubules. Biophysical Journal, 2012. 102(6): p. 1274–1283.

29. Janosi, I.M., D. Chretien, and H. Flyvbjerg, Modeling elastic properties of microtubule tips and walls. European Biophysics Journal with Biophysics Letters, 1998. 27(5): p. 501–513.

30. Molodtsov, M.I., Ermakova, E. A., Shnol, E. E., Grishchuk, E. L., McIntosh, J. R., Ataullakhanov, F. I., A molecular-mechanical model of the microtubule. Biophys. J., 2005. 88(5): p. 3167–3179.

31. VanBuren, V., Cassimeris, L., Odde, D. J., Mechanochemical model of microtubule structure and self-assembly kinetics Biophys. J., 2005. 89(5): p. 2911–2926.

32. Zong, C.H., Lu, T., Shen, T. Y. and Wolynes, P. G., Nonequilibrium self-assembly of linear fibers: microscopic treatment of growth, decay, catastrophe and rescue. Phys. Biol., 2006. 3(1): p. 83–92.

33. Dimitrov, A., et al., Detection of GTP-Tubulin Conformation in Vivo Reveals a Role for GTP Remnants in Microtubule Rescues. Science, 2008. 322(5906): p. 1353–1356.

34. Gillespie, D.T., General Method for Numerically Simulating Stochastic Time Evolution of Coupled Chemical-Reactions. Journal of Computational Physics, 1976. 22(4): p. 403–434.

35. Gillespie, D.T., Stochastic simulation of chemical kinetics. Annual Review of Physical Chemistry, 2007. 58: p. 35–55.

36. Alushin, G.M., et al., High-Resolution Microtubule Structures Reveal the Structural Transitions in alpha beta-Tubulin upon GTP Hydrolysis. Cell, 2014. 157(5): p. 1117–1129.

37. Perutz, M.F., Stereochemistry of Cooperative Effects in Haemoglobin. Nature, 1970. 228(5273): p. 726-&.

38. Frauenfelder, H., S.G. Sligar, and P.G. Wolynes, The Energy Landscapes and Motions of Proteins. Science, 1991. 254(5038): p. 1598–1603.

39. Driver, J.W., et al., Direct measurement of conformational strain energy in protofilaments curling outward from disassembling microtubule tips. Elife, 2017. 6.

40. Voter, W.A., E.T. Obrien, and H.P. Erickson, Dilution-Induced Disassembly of Microtubules - Relation to Dynamic Instability and the Gtp Cap. Cell Motility and the Cytoskeleton, 1991. 18(1): p. 55–62.

41. Walker, R.A., N.K. Pryer, and E.D. Salmon, Dilution of Individual Microtubules Observed in Real-Time Invitro - Evidence That Cap Size Is Small and Independent of Elongation Rate. Journal of Cell Biology, 1991. 114(1): p. 73–81.

42. Coombes, C.E., et al., Evolving Tip Structures Can Explain Age-Dependent Microtubule Catastrophe. Current Biology, 2013. 23(14): p. 1342–1348.

43. Zakharov, P., et al., Molecular and Mechanical Causes of Microtubule Catastrophe and Aging. Biophysical Journal, 2015. 109(12): p. 2574–2591.

44. Rice, J.A., Mathematical Statistics and Data Analysis1995, Belmont, CA: Duxbury Press.

45. Caplow, M. and J. Shanks, Evidence that a single monolayer tubulin-GTP cap is both necessary and sufficient to stabilize microtubules. Molecular Biology of the Cell, 1996. 7(4): p. 663–675.

46. Akhmanova, A. and M.O. Steinmetz, Tracking the ends: a dynamic protein network controls the fate of microtubule tips. Nature Reviews Molecular Cell Biology, 2008. 9(4): p. 309–322.

47. Fernandez, N., Chang, Q., Buster, D. W., Sharp, D. J., Ma, A., A model for the regulatory network controlling the dynamics of kinetochore microtubule plus-ends and poleward flux in metaphase. Proceedings of the National Academy of Sciences of the United States of America, 2009. 106(19): p. 7846–7851.

48. Goodwin, S.S. and R.D. Vale, Patronin Regulates the Microtubule Network by Protecting Microtubule Minus Ends. Cell, 2010. 143(2): p. 263–274.

49. Tanaka, N., et al., Nezha/CAMSAP3 and CAMSAP2 cooperate in epithelial-specific organization of noncentrosomal microtubules. Proceedings of the National Academy of Sciences of the United States of America, 2012. 109(49): p. 20029–20034.

50. Jiang, K., et al., Microtubule Minus-End Stabilization by Polymerization-Driven CAMSAP Deposition. Developmental Cell, 2014. 28(3): p. 295–309.

51. Currie, J.D., et al., The microtubule lattice and plus-end association of Drosophila Mini spindles is spatially regulated to fine-tune microtubule dynamics. Molecular Biology of the Cell, 2011. 22(22): p. 4343–4361.

52. Vitre, B., et al., EB1 regulates microtubule dynamics and tubulin sheet closure in vitro. Nature Cell Biology, 2008. 10(4): p. 415-U81.

53. Al-Bassam, J., et al., CLASP Promotes Microtubule Rescue by Recruiting Tubulin Dimers to the Microtubule. Developmental Cell, 2010. 19(2): p. 245–258.

54. Serebrinsky, S.A., Physical time scale in kinetic Monte Carlo simulations of continuous-time Markov chains. Physical Review E, 2011. 83(3).

